# Targeted delivery of a phosphoinositide 3-kinase γ inhibitor to restore organ function in sepsis through dye-functionalized lipid nanocarriers

**DOI:** 10.1101/2021.01.20.427305

**Authors:** Adrian T. Press, Petra Babic, Bianca Hoffmann, Tina Müller, Wanling Foo, Walter Hauswald, Jovana Benecke, Martina Beretta, Zoltán Cseresnyés, Stephanie Hoeppener, Ivo Nischang, Sina M. Coldewey, Markus H. Gräler, Reinhard Bauer, Falk Gonnert, Nikolaus Gaßler, Reinhard Wetzker, Marc Thilo Figge, Ulrich S. Schubert, Michael Bauer

## Abstract

Jaundice, the clinical hallmark of infection-associated liver dysfunction, reflects altered membrane organization of the canalicular pole of hepatocytes and portends poor outcomes. Mice lacking phosphoinositide 3-kinase-γ (PI3Kγ) are protected against membrane disintegration and hepatic excretory dysfunction. However, they exhibit a severe immune defect that hinders neutrophil recruitment to sites of infection. To exploit the therapeutic potential of PI3Kγ inhibition in sepsis, a targeted approach to deliver drugs to hepatic parenchymal cells without compromising other cells, in particular immune cells, seems warranted. Here we demonstrate that nanocarriers functionalized through DY-635, a fluorescent polymethine dye and a ligand of organic anion transporters can selectively deliver therapeutics to hepatic parenchymal cells. Applying this strategy to a murine model of sepsis, we observed PI3Kγ-dependent restoration of biliary canalicular architecture, maintained excretory liver function, and improved survival without impairing host defense mechanisms. This strategy carries the potential to expand targeted nanomedicines to disease entities with systemic inflammation and concomitantly impaired barrier functionality.

**One-Sentence Summary:** Dye-functionalized liposomes allow delivery of a PI3Kγ inhibitor to hepatocytes to resolve sepsis-related liver failure without ‘off-target’ effects on immunity.

**Graphical Abstract:** Targeting PI3Kγ in hepatocytes by dye-functionalized liposomes to resolve sepsis-related liver failure without ‘off-target’ effects on immunity.

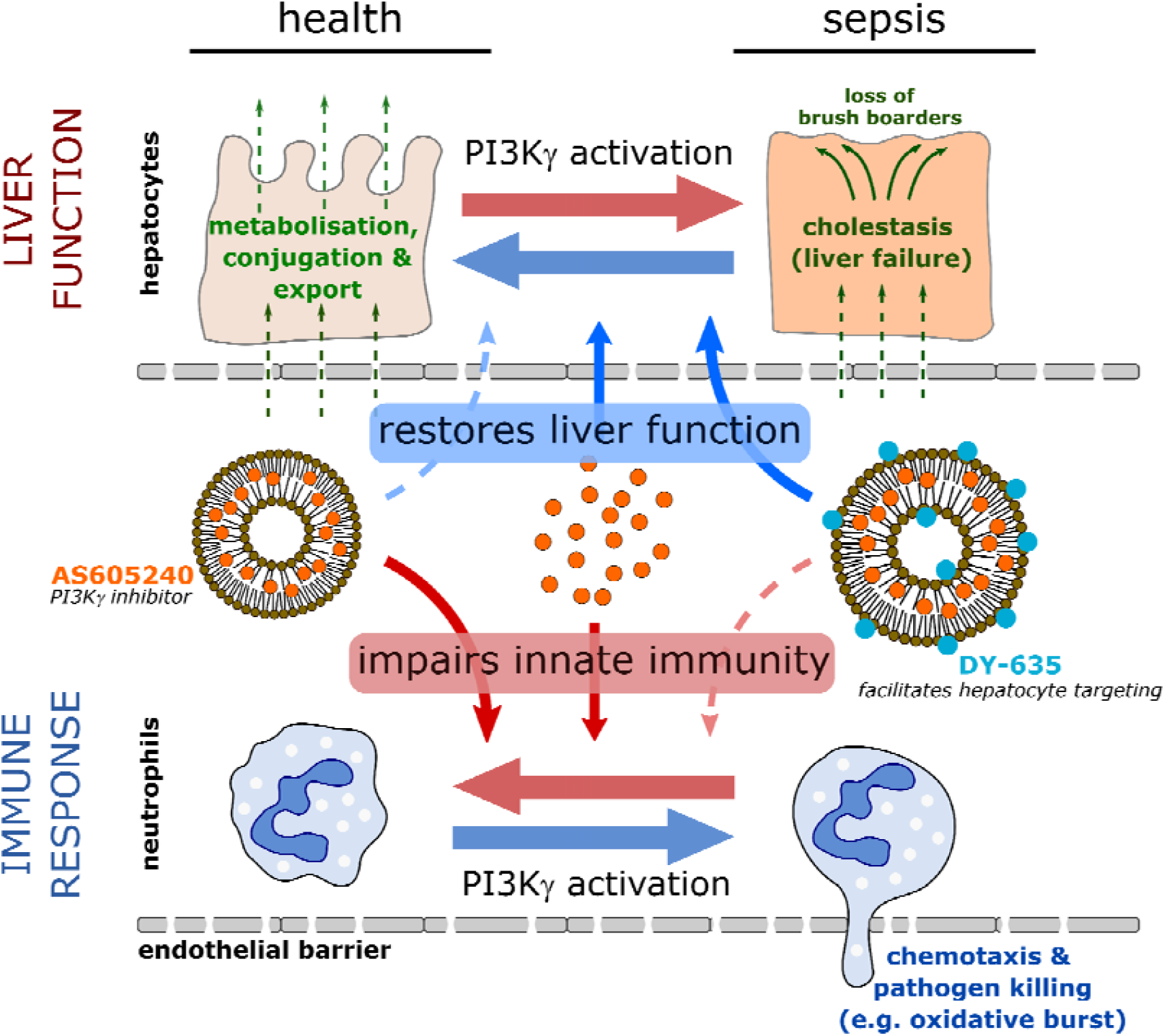

## Main

Sepsis remains a major cause of death despite advances in medical care. The current ‘Sepsis-3’ definition shifted the conceptual focus away from an exaggerated systemic inflammatory response towards an abreaction by the host’s defense mechanisms affecting both directly involved and remote organs.^1^ In this regard, metabolic (mal)adaptation appears to be of crucial significance.^2,3^

We previously demonstrated that mice lacking PI3Kγ were protected against hepatic excretory dysfunction (i.e. disturbed phase I and II metabolism and transcellular transport) during sepsis^4^ yet a concurrent and severe immune defect rendered them vulnerable to peritonitis. This presumably reflects the key role of PI3Kγ in neutrophil migration.^5^ Thus, conflicting functions of PI3Kγ in liver parenchyma and in immune cells regarding net outcome of inhibition might be hypothesized. Expression of the PI3Kγ gene by cells other than neutrophils has received some attention^6^ but, to our knowledge, has not yet been explored in hepatocytes. Unequivocally, the ability to successfully target PI3Kγ to specific (parenchymal) cells while avoiding potentially deleterious effects on immune function would offer a fundamental improvement in therapeutic interventions for sepsis, a condition where multiple adjunct therapies have failed over decades.^7,8^

Nanomedicine holds great promise but requires design of therapeutics with a distinct combination of features such as low toxicity, simplicity and efficacy.^9^ The pharmacology and toxicology of nanocarriers depend on size, composition, surface charge and shape; these all determine uptake, immune cell recognition, and circulation time. Conversely, an enhanced permeability and retention (EPR) effect leads to passive enrichment of the cargo-carrier systems in tissues where the barrier function is locally altered (e.g. by tumor or local inflammation), reflecting a significant hurdle for conditions such as sepsis, severe tissue trauma or hemorrhagic shock where barriers are ubiquitously disturbed.^10,11^ Modifying nanocarriers with dyes known to act as ligands of organic anion transporters may enable substantial enrichment of the drug cargo into hepatocytes while, at the same time, reducing uptake by circulating neutrophils or tissue macrophages (e.g. liver Kupffer cells) lacking these transporters.^12^

In the current investigation, we aimed to restore liver excretory function by local targeting of PI3Kγ. We hypothesized that selective hepatocellular targeting of PI3Kγ attenuates liver failure in peritonitis-induced sepsis while avoiding immune functionality and untoward side effects on the peritoneal host response. To test this postulate, we used dye-functionalized liposomes to deliver AS605240, a PI3Kγ inhibitor, to hepatic parenchymal cells in a model of severe polymicrobial peritonitis where side effects compromising immunity would be clearly unwelcome.

## Results

### Expression of PI3Kγ in human liver and effect of PI3Kγ null mice on sepsis survival

To scrutinize the cell-type-specific involvement of PI3Kγ in sepsis-induced hepatic excretory dysfunction and to characterize the protein as a therapeutic target, expression of the signaling protein in murine and human liver tissue was first investigated. We used a validated polyclonal antibody raised from an orthologue peptide sequence of PI3Kγ in mouse and humans.^13^ Liver tissue slices were prepared from nine patients with very mild to severe liver pathologies including non-alcoholic fatty liver disease, liver cirrhosis and acute liver failure (**Supplementary Table 1**).

In all tissue sections, we found consistent staining of PI3Kγ in hepatocytes, bile ducts and larger vessels. The intensity of PI3Kγ expression was expectedly high in infiltrating immune cells, primarily neutrophils occasionally found within the sections (**Figure 1A**). Other non-parenchymal liver cells, in particular Kupffer cells and liver sinusoidal endothelial cells were negative for PI3Kγ. To validate these results, we performed Western blots with the same antibody against PI3Kγ in a sample of human non-parenchymal cells (NPCs) and primary human hepatocytes obtained from a pool of 20 male or 20 female donors. In line with the immunohistochemistry staining of patient biopsies, Western blots revealed expression of PI3Kγ in primary human hepatocytes and leukocytes obtained from healthy volunteers, while liver non-parenchymal cells, i.e. sinusoidal endothelial cells, Kupffer cells and cholangiocytes were negative for the signaling protein (**Figure 1B**). Expression of PI3Kγ was similarly observed in murine hepatocytes and further enhanced in primary human hepatocytes and murine hepatocytes after stimulation with a mix of pro-inflammatory cytokines (LPS, IFN-γ, IL-1β and TNF-α) for 24 h. This indicates involvement of the signaling protein within the inflammatory response of these cells (**Figure 1C**).

**Figure 1.**
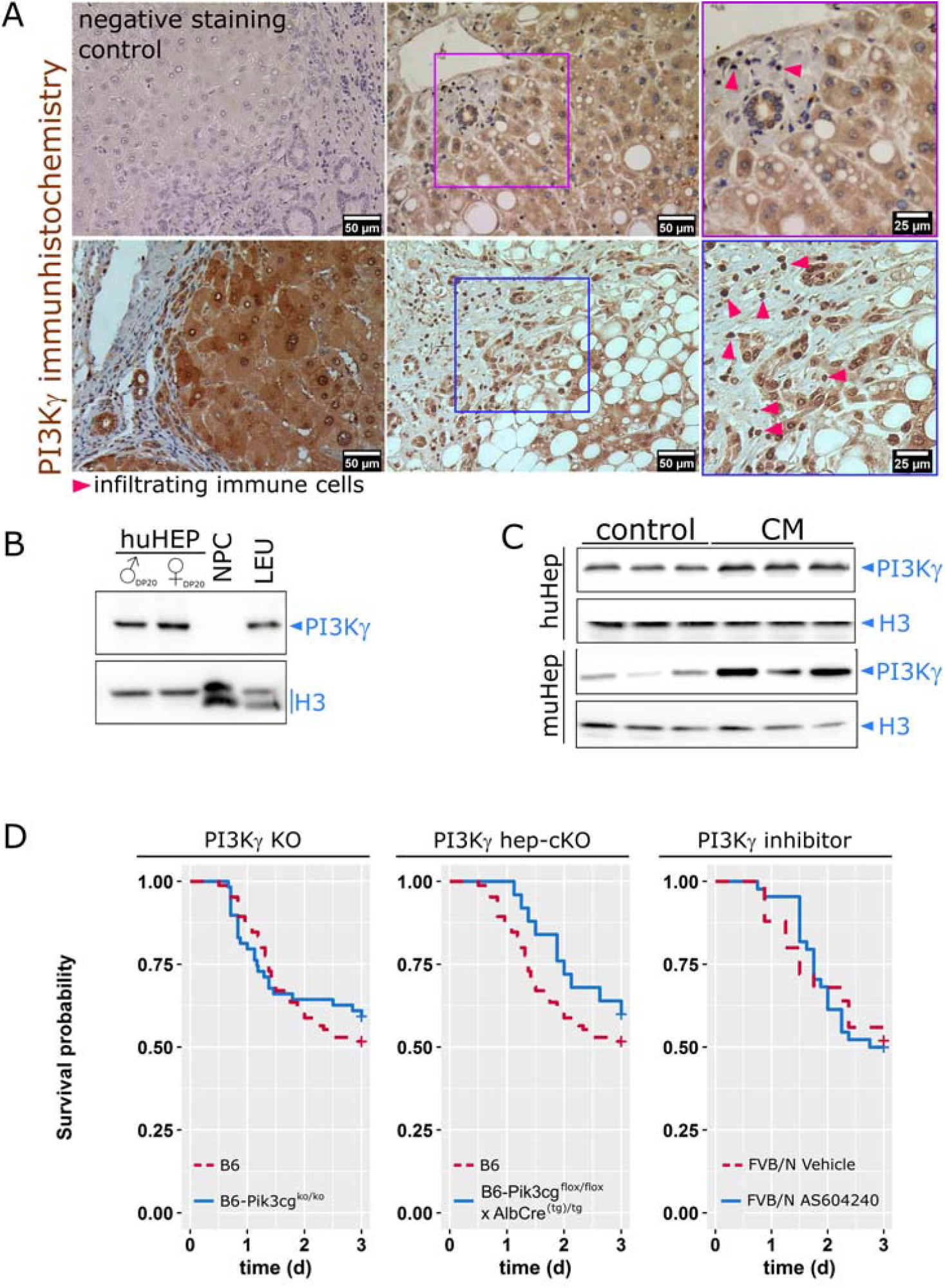
Cell-type specific expression pattern of PI3Kγ in the liver, induction and function under conditions of inflammation. **(A)** PI3Kγ is expressed by human hepatocytes and infiltrating immune cells in biopsies from patients with minimal to mild inflammatory activity. Triangles point to immune cell (clusters) including some neutrophils which are known to highly express PI3Kγ. In the negative control the primary antibody was replaced by an equal volume buffer. **(B)** PI3Kγ expression in human primary hepatocytes from 20 male (♂) or female (♀) donor pools (HEP, DP20), but not non-parenchymal cells (NPC). Isolated human leukocytes from healthy volunteers (LEU) served as positive controls. **(C)** Primary murine hepatocytes and HepG2 cells expressed PI3Kγ under basal conditions; expression was increased at 24 h after stimulation with LPS, IFN-γ, IL-1β and TNF-α (CM).**(D)** Survival of WT, PI3Kγ null (left) and liver specific PI3Kγ knockout mice (PI3Kγ flox^tg/tg^ x AlbCre^(tg)/tg^, middle) or systemic application of the PI3Kγ inhibitor, AS605240 (right) in a model of polymicrobial sepsis induced by peritoneal contamination with a human stool suspension.

We next investigated the effect of systemic and tissue specific knock-out of PI3Kγ on the survival in our murine sepsis model. As shown in **Figure 1D** no significant difference in survival rates of wild-type and PI3Kγ knock-out mice entities was observed, suggesting balanced beneficial and detrimental effects of PI3Kγ deficiency upon the host response. Septic PI3Kγ knockout mice did not develop excretory liver failure^4^ whereas the liver-specific, but not the systemic PI3Kγ knockout treatment resulted in a trend towards increased survival in the initial phase of sepsis (**Figure 1D**). Knock-out of PI3Kγ deleted kinase activity as well as kinase-independent functions that, for example, contribute to integrity of hepatocyte canalicular membrane.^14^ Together these findings motivated us to attempt hepatocyte-specific pharmacological selective targeting of the kinase function of PI3Kγ using a small molecular inhibitor of kinase activity as a therapeutic concept.

### Characterization and pharmacological properties of dye-functionalized lipid nanoparticles for hepatocellular drug delivery

The thiazolidine compound AS605240 was selected as a specific inhibitor of PI3Kγ enzymatic activity, that is known to inhibit neutrophil migration and function in a dose-dependent manner.^15,16^ To specifically direct the drug to liver parenchyma, we used liposomes containing 1,2-dipalmitoyl-sn-glycerol-3-phosphoethanolamine (DPPE) azide modified with the fluorescent dye DY-635. DY-635 was recently characterized as a molecule specifically recognized and eliminated by hepatocytes.^12^ DY-635 conjugated with DPPE was then used to prepare targeted liposomes that deliver the PI3Kγ inhibitor AS605240 into hepatocytes. The liposomes used in this study (five batches) contained 4.3 ± 1.71 µmol L^−1^ AS605240 and 130 ± 30 pmol L^−1^ DY-635 (mean ± SD) (**Figure 2A**). All liposomes revealed long-term stability with mean hydrodynamic diameters (dynamic light scattering (DLS), intensity mode) of 150-160 nm and polydispersity index (PDI) of 0.2. Only non-targeted, AS605240-loaded liposomes (NT-LipoAS) were smaller (average ∼90 nm) and had a lower PDI of 0.1. (**Figure 2B**) The determination of the diameter from cryo-transmission electron microscopy (cryo-TEM) images from different liposomal preparations confirmed the hydrodynamic diameters from the DLS analysis. For the AS605240-loaded liposomes (T-LipoAS and NT-LipoAS), as well as the non-targeted AS605240-free liposomes, the diameter was smaller compared to the hydrodynamic diameter obtained from DLS. This difference could have resulted from variations in the hydration of the different carriers. The DLS of the drug-free carrier, T-Lipo revealed a slightly higher diameter (190 nm). Quantification of the diameters by transmission electron microscopy confirmed the measurements obtained by DLS (**Figure S1**). Liposomes were further characterized by analytical ultracentrifugation. Liposomes did not sediment at a measured fluid density (□_0_) of 1.0822 g cm^−1^ and dynamic viscosity (η_0_) of 1.1996 mPas of a heavy water/buffer mixture, suggesting a rather high partial specific volume (ν) of ∼0.924 cm^3^ g^−1^ (**Figure S2**). The sedimentation coefficient distributions suggest a hydrodynamic diameter of ∼ 150 nm (for T-Lipo) and 141 nm (for T-LipoAS).^17^ These values are slightly smaller than the hydrodynamic diameters obtained by DLS. Liposomes diluted in the buffer showed stability during sedimentation at 10 000 rpm and re-dispersibility by simple shaking of the sedimented material in the centrifuge cells.

**Figure 2.**
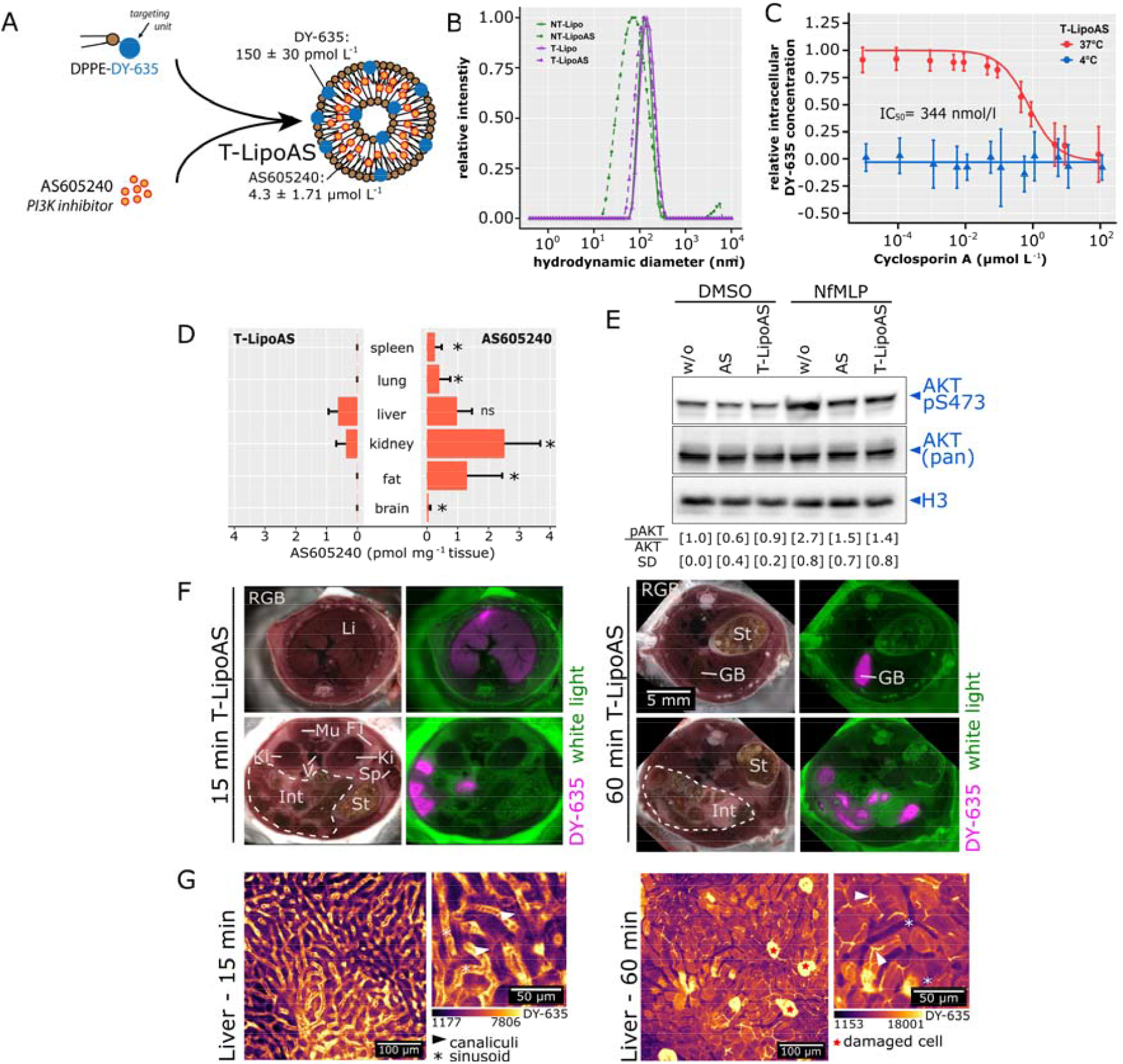
Formulation of targeted liposomes, *in vitro* uptake characteristics, use for drug delivery and pharmacodynamic properties. **(A)** Scheme for the preparation of DY-635 targeted liposomes containing the PI3Kγ inhibitor AS605240 (T-LipoAS). The concentrations of AS605240 and DY-635 were determined by LC-MS from five individually prepared batches used for all subsequent experiments. **(B)** Hydrodynamic diameters of different liposomes in histidine buffer analyzed by dynamic light scattering. **(C)** Uptake characteristics of T-LipoAS by primary human hepatocytes demonstrates competitive inhibition by ciclosporin A- and energy dependence, confirming active uptake by organic anion transporters (OATPs). (**D**) Quantification of AS60524060 min after injection of 1 mg kg^−1^ AS605240 (i.p. in DMSO) or 2.5 mg kg^−1^ T-LipoAS (i.v.) in tissue lysates by LC-MS. *p<0.05 for Wilcoxon test between T-LipoAS and AS605240 in each individual tissue. (**E**) T-LipoAS effectively inhibits fMLP-stimulated PI3Kγ activity, reflected by inhibition of AKT S473 phosphorylation. Experiments were conducted with hepatocytes from three different donors. The mean pAKT/AKT ratio and standard deviation (SD) were calculated for all experiments separately. **(F)** Episcopic imaging of mice treated with T-LipoAS for 15 or 60 min and relative concentration of the targeting moiety DY-635 in the different compartments. (**G**) Intravital microscopy to monitor T-LipoAS kinetics in the liver. Liposomes appear in the vasculature after 5-15 minutes and start to selectively accumulate within hepatocytes. After 35-60 min the bile canalicular system appears stained, indicating the processing and elimination of the targeting moiety (DY-635) from T-LipoAS.

DY-635 facilitates targeted delivery of nanocarriers to hepatocytes and specific tumor cells expressing organic anion transporters and pumps (OATs and OATPs).^12,18,19^ To determine whether the targeting unit on liposomes improves hepatocyte uptake, we used transporter-qualified primary human hepatocytes expressing various OATs and OATPs among other basolateral hepatocellular uptake transporters. Incubation with T-LipoAS at 37°C led to a strong uptake that was diminished by ciclosporin A, which acts as an competitive inhibitor OAT(P)s. The calculated inhibitory concentration (IC)_50_ of 344 nmol L^−1^ was similar to a previously reported value for DY-635-conjugated polymer nanoparticles.^12^ The ciclosporin A-sensitive uptake was quenched by lowering temperature to 4°C, indicating the active and OAT(P) mediated uptake-process (**Figure 2C**).

Next, we analyzed tissue distribution of AS605240 in mice at 60 min post-injection of free AS605240 (4 mg kg^−1^, intraperitoneal) or T-LipoAS (2.5 mg kg^−1^, intravenous). The doses were matched based on preliminary experiments to achieve comparable concentrations within the liver. Free AS605240 injected intraperitoneally accumulated in all analyzed tissues (spleen, lung, liver, kidney, fat and brain), while AS605240 derived from T-LipoAS was only found in liver and kidney. The concentrations of AS605240 accumulated in the liver at 60 min after injection did not significantly differ between formulations (**Figure 2D**), confirming enriched accumulation of the compound in the liver while avoiding other organs except the kidney. To elaborate the inhibitory capacity of AS605240 after nanoformulation, its inhibitory effects on PI3Kγ-mediated AKT serine (S)473 phosphorylation induced by N-formylmethionine-leucyl-phenylalanine (fMLP) was investigated in primary human hepatocytes confirming functional delivery by the liposomal formulation (**Figure 2E)**.

This notion was confirmed by episcopic imaging detecting DY-635 fluorescence from injected T-LipoAS to investigate the biodistribution of the carrier. As early as 15 min post-administration of T-LipoAS, there was a marked increase in DY-635 signal intensity in liver tissue and, after hepatobiliary excretion, in proximal parts of the small intestine. By contrast, little to no DY-635 fluorescence signal was contributed by spleen or lung. After 1 h DY-635 was mainly eliminated through the hepatobiliary system as indicated by the stained gall bladder and gastrointestinal lumen (**Figure 2F**). The liver consists of various cell types controlling metabolism and the immune response. Kupffer cells (the liver macrophage population) and liver-specific endothelial cells (LSECs) have been shown to sequester nanocarriers.^12,20,21^ Both cell types would be considered “off-target” for our delivery strategy since they strongly contribute to the immune response against invading microorganisms. Therefore cell-type specific targeting of loaded liposomes within the liver was further investigated by time-lapse intravital microscopy (**Figure 2G**). This method enables differentiation of parenchymal (hepatocytes) and non-parenchymal (e.g. Kupffer cells, LSECs) within the liver due to their different shape, location and NAD(P)H fluorescence. After intravenous injection, T-LipoAS started to appear in the capillary system (sinusoids) (**Figure 2G**). T-LipoAS then rapidly accumulated within hepatocytes. Clearance of nanocarriers by Kupffer cells or LSECs can be identified with this method via monitoring of line-patterns or spots at the interphase between NADPH-positive hepatocytes and the sinusoids.^9,17^ No signal of T-LipoAS accumulated at these locations. The appearance of line structures within the hepatocytes indicate the accumulation of DY-635 in canaliculi, the smallest structure formed by hepatocytes within the bile drainage pathway,further confirming the hepatobiliary elimination of DY-635. The apparent lack of non-parenchymal cells in the liver (e.g. Kupffer cells or LSECs) in combination with the appearance of DY-635 derived from T-LipoAS within hepatocytes and canaliculi indicates a highly selective uptake and processing of targeted nanocarriers through hepatocytes.

Cell culture models cannot resemble the complexity of the pathophysiological mechanisms of sepsis, and cannot provide evidence that the suggested targeting properties of T-LipoAS are maintained under such circumstances. Therefore, the targeting procedure was tested in a complex rodent model of sepsis based on peritoneal contamination and infection (PCI). In this model human feces are administered intraperitoneally to induce life-threatening polymicrobial infection. This model is well characterized with signs of excretory liver failure detectable as early as 6 h after initiation of infection.^4,22–24^ To explore the targeting approach animals were treated with T-LipoAS at 6 h after infection with liver function assessed at 24 h. All animals received supportive antibiotic (meropenem) and analgesic (metamizole) therapy (**Figure 3A)**.

**Figure 3.**
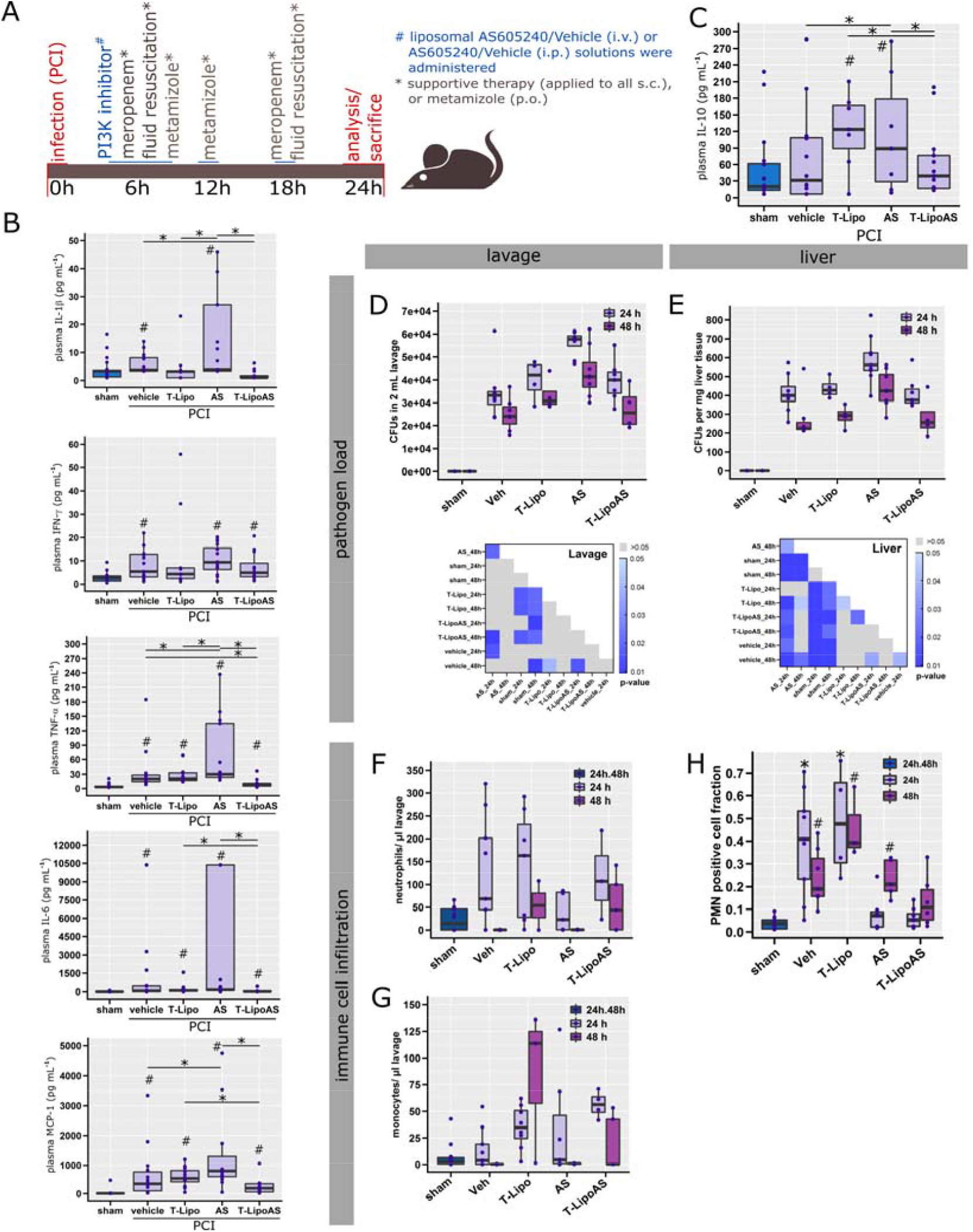
Targeted delivery of AS605240 prevents untoward effects on the innate immune response to peritonitis. **(A)** Graphical summary of the murine sepsis model and experimental design. **(B)** Pro- and **(C)** anti-inflammatory cytokines were analyzed in EDTA-plasma from peritoneal contamination and infection (PCI) and sham animals. ^#^p<0.05 against sham, *p<0.05 as indicated; Kruskal-Wallis ANOVA with controlled false-discovery rates (Benjamini-Hochberg procedure). **(D,E)** Colony forming units (CFUs) of bacteria reflecting host defense failure were analyzed from (**D**) abdominal lavage and (**E**) liver homogenates. Significance maps depict p-values from a Kruskal-Wallis ANOVA with controlled false-discovery rates applying the Benjamini-Hochberg procedure. **(F,G)** Concentration of (**F**) neutrophils and (**G**) monocytes were quantified in 2 mL of peritoneal lavage. (**H**) Infiltrating polymorphonuclear neutrophils (PMN) were assessed from histological sections after immune fluorescent staining. *p<0.05 against sham, ^#^p<0.05 against 24h; Kruskal-Wallis ANOVA with controlled false-discovery rates (Benjamini-Hochberg procedure).

PI3Kγ knock-out mice display a “cytokine storm” in response to severe infection that exceed levels seen in wild-type animals.^4^ Similar effects were observed for non-targeted pharmacologic inhibition of PI3Kγ in the present study. Several pro-inflammatory cytokines (IL-1α, IL-1β, IL-6, IL-27, IFN-β, IFN-γ, TNF-α, GM-CSF, and MCP-1) and the anti-inflammatory cytokine, IL-10 were analyzed. In septic animals treated with free AS605240 (PCI, AS group) IFN-β, TNF-α, GMCSF, MCP-1, and IL-10 levels were significantly elevated in comparison to septic animals treated with vehicle (PCI, Vehicle group). IL-1α, IL-1β, IFN-γ, IL-27 trended towards an increase **(Figure 3B,C, Figure S3A)**. An unsupervised clustering of the animals, performed according to similarities in their cytokine profile, also suggested a ‘cytokine storm’ was present, particularly in septic animals treated with free AS605240 **(Figure S3B)**. This untoward effect on systemic inflammation was not observed when the inhibitor was formulated within liposomes (T-LipoAS) (**Figure 3B, Figure S3A,B**). In contrast to the obvious pro-inflammatory effect of free AS605240, the absence of the cytokine storm in T-LipoAS-treated animals is consistent with specific targeting of the PI3Kγ inhibitor in parenchymal liver cells but immune cell compartment sparing (**Figure 2C**).

To further investigate this critical aspect of cell-type specificity of T-LipoAS, we examined innate immunity (bacterial clearance and neutrophil recruitment) using functional assays. Most importantly, intraperitoneal-administered AS605240 in the PCI-model resulted in an increased pathogen load within the peritoneal cavity (**Figure 3D**), the primary site of infection, and in liver tissue (**Figure 3E**). This was absent when hepatocellular inhibition of PI3kγ was achieved by T-LipoAS. Thus, the increased pathogen load may reflect off-target effects of impaired immune cell recruitment and function after intraperitoneal administration of AS605240 to achieve hepatocellular PI3kγ inhibition. Consequently, lower concentrations of neutrophils (**Figure 3F**) and monocytes (**Figure 3G**) were detected within the peritoneal cavity.

The number of neutrophils infiltrating into liver tissue was increased in untreated animals with sepsis (PCI, vehicle group) (**Figure 3H**). By contrast, hepatic parenchymal infiltration of neutrophils in both AS and T-LipoAS treated septic animals (PCI, AS or T-LipoAS group) was significantly reduced. As the T-LipoAS treated mice express low levels of circulating cytokines, this result may reflect a secondary effect of a reduced local inflammatory response (**Figure 3B,C**) due, for example, to te protection of liver function (**Figure 4**), rather than the result of systemic immunosuppression. In support of this assumption, no difference in vehicle and T-LipoAS treated animals were observed with respect to the pathogen load in the liver (**Figure 3E**). This hypothesis is further strengthened by the results of neutrophils infiltration pattern into the peritoneal cavity upon infection. While animals treated with AS605240 showed significantly lowered numbers of peritoneal neutrophils, the numbers of neutrophils accumulating in the peritoneal cavity upon infection when PI3Kγ was selectively targeted with T-LipoAS were similar to septic animals treated with vehicle.

**Figure 4.**
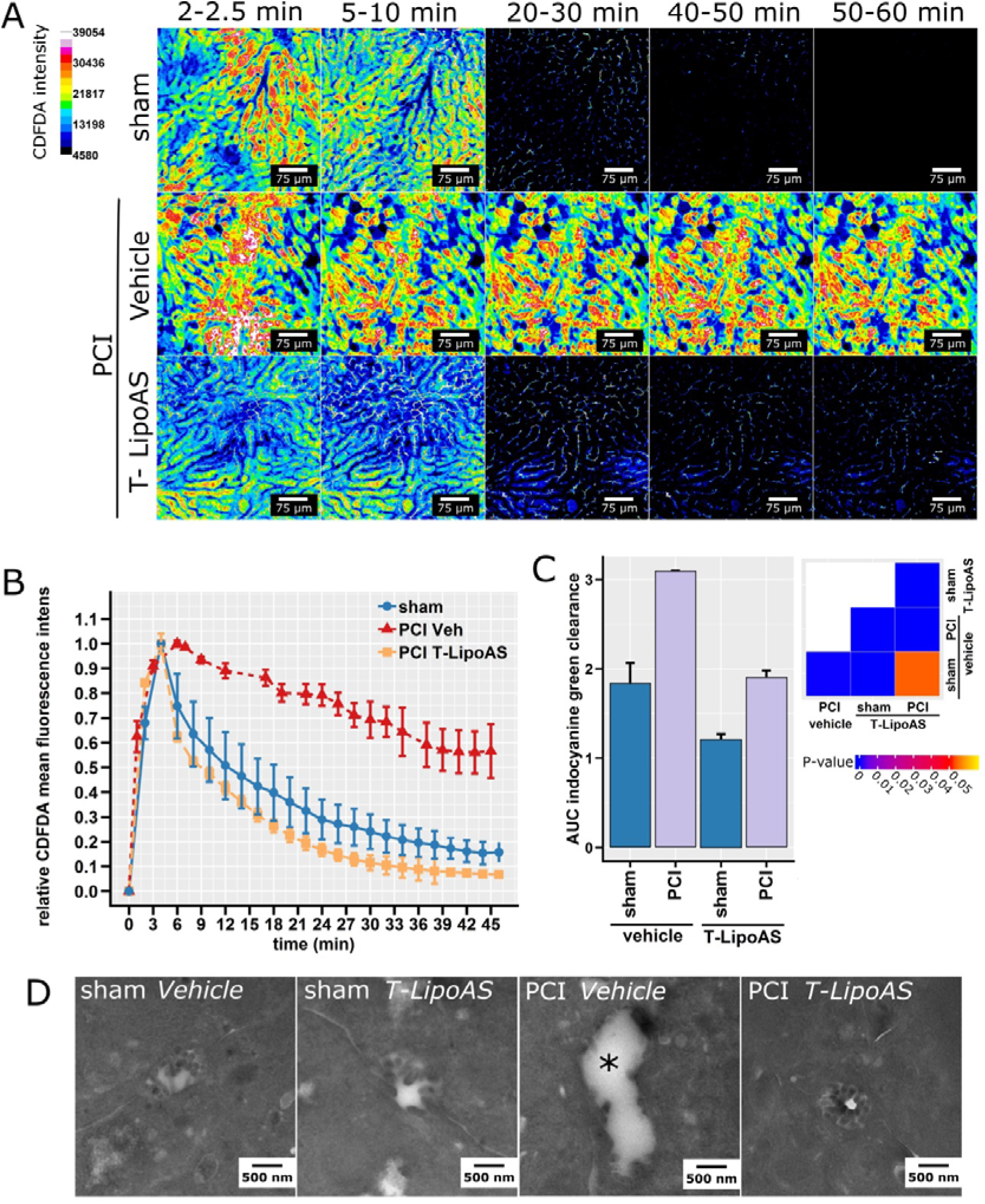
Nanoformulated PI3Kγ inhibition restores excretory liver function *in vivo*. **(A)** Intravital microscopy of elimination of CDFDA, a fluorescent substrate subject to excretory elimination by hepatocytes via Mrp-2, allows direct visualization of excretory function in sham or septic (PCI) animals treated with vehicle or a hepatocyte-directed inhibitor of PI3Kγ (T-LipoAS). **(B)** Normalized CDFDA intensities quantified from intravital images depict plasma disappearance curves (mean ± SD). Sham and PCI animals treated with T-LipoAS both showed a classical biphasic decay curve while PCI vehicles showed a rather linear decay and a higher area under the curve indicative of excretory liver failure. **(C)** Similar results were obtained when liver excretory function was tested by indocyanine green clearance. Indocyanine green was detected non-invasively by multispectral optoacoustic tomography. The ability to clear ICG was quantitatively analyzed by comparing areas under the curve (AUC). **(D)** Representative transmission electron micrographs from bile canaliculi of healthy control (sham) and septic animals (PCI) treated with vehicle or the PI3Kγ inhibitors, free AS or T-LipoAS. The asterisk marks a bile canaliculus where typical brush borders are lost due to sepsis.

As mentioned above, and based on previous observations in PI3Kγ knockout mice,^4^ we hypothesized that the protective effect of hepatocyte-targeted PI3Kγ-inhibition was due to preservation of excretory liver function. Three different approaches were used to investigate the effects of T-LipoAS on excretory liver failure after 24 h of sepsis (**Figure 3A**). First, intravital microscopy was applied to assess accumulation and clearance of the Mrp2 substrate, CDFDA as a surrogate for excretory function of endogenous substrates of this pathway, e.g. bilirubin (**Figure 4 A,B**). CDFDA in control (sham animals, injected i.p. with sterile saline) accumulated rapidly in the liver, specifically in hepatocytes, but was effectively eliminated within 60 min. In sepsis (PCI) however, CDFDA accumulated as rapidly as in the reference group but elimination was visibly prolonged, as reflected by a change in elimination kinetics. Intriguingly, treatment with T-LipoAS fully resolved the effects of sepsis on liver excretory function; the initial accumulation and elimination was similar to that seen in non-infected sham animals (**Figure 4B**). To further validate these findings, we investigated the plasma disappearance rate of indocyanine green (ICG), a clinical approved dye used to assess excretory liver function. Plasma disappearance is analyzed in patients by pulse densitometry but such devices are lacking for use in small rodents. Due to the (near)infrared (NIR) absorbance of ICG above 750 nm, multispectral optoacoustic tomography can be employed, allowing non-invasive detection of (N)IR at up to video frame rate and with good spatial resolution. Applying automated image analysis algorithms, we identified suitable areas for quantification of ICG clearance over time in each animal and investigated elimination through quantification of area under the curve (AUC) (**Figure S4**). The increased AUC reflects decreased elimination of ICG from plasma and correlates with increased hepatic ICG retention, i.e. lowered elimination capability. Similar to CDFDA clearance, ICG also accumulated significantly in animals at 24 h post-induction of sepsis (**Figure 3C**). This effect was also prevented by a single injection of liposomal AS605240 (T-LipoAS) for hepatocyte-targeted PI3Kγ inhibition at 6 h.

The loss of the ability of hepatocytes to eliminate endo- and xenobiotics, as probed by the dye-clearance assays, was reflected by structural changes such as the loss of canalicular brush borders. In line with previous reports, the majority of canalicular brush borders was lost at 24 h, while these structures remained intact in mice treated with the nanoformulated PI3Kγ inhibitor AS605240 (T-LipoAS), similar to our previous observation in PI3Kγ knockout mice (**Figure 4D**).^4^ Taken together, these findings demonstrate morphological and functional protection of hepatocytes by treatment with the PI3K inhibitor AS605240 in a hepatocyte-targeted carrier.

Since sepsis-associated liver failure is linked with poor survival,^25,26^ targeted delivery of AS605240 through T-LipoAS improved not only liver function but also overall outcome (**Figure 5**). The chosen sepsis model aimed to mimic a clinical situation. Animals were infected with a large dose of fecal slurry followed by administration of meropenem, a broad-spectrum antibiotic, to achieve an approximate 50% lethality. In the absence of any surgical procedure to clear bacteria from the abdominal cavity (the primary site of infection) chronic abscesses are formed making survival analysis beyond seven days difficult to interpret. Different AS605240 and vehicle formulations were administered daily over the first five days, commencing at 6 h post-infection (**Figure 5A**). No survival difference was observed between animals treated systemically with AS605240 and septic controls (**Figure 5B**). AS605240 formulated in non-targeted liposomes (NT-LipoAS) even worsened mortality, perhaps due to improved bioavailability and immune-cell clearing of non-targeted liposomes compared to the targeted formulations (**Figure S5**). By contrast, the DY-635 formulation, T-LipoAS both prolonged and significantly increased survival (**Figure 5C, Figure S5**), supporting findings that the hepatocyte-specific delivery of AS605240 was achieved and off-target effects on the immune system significantly reduced.

**Figure 5.**
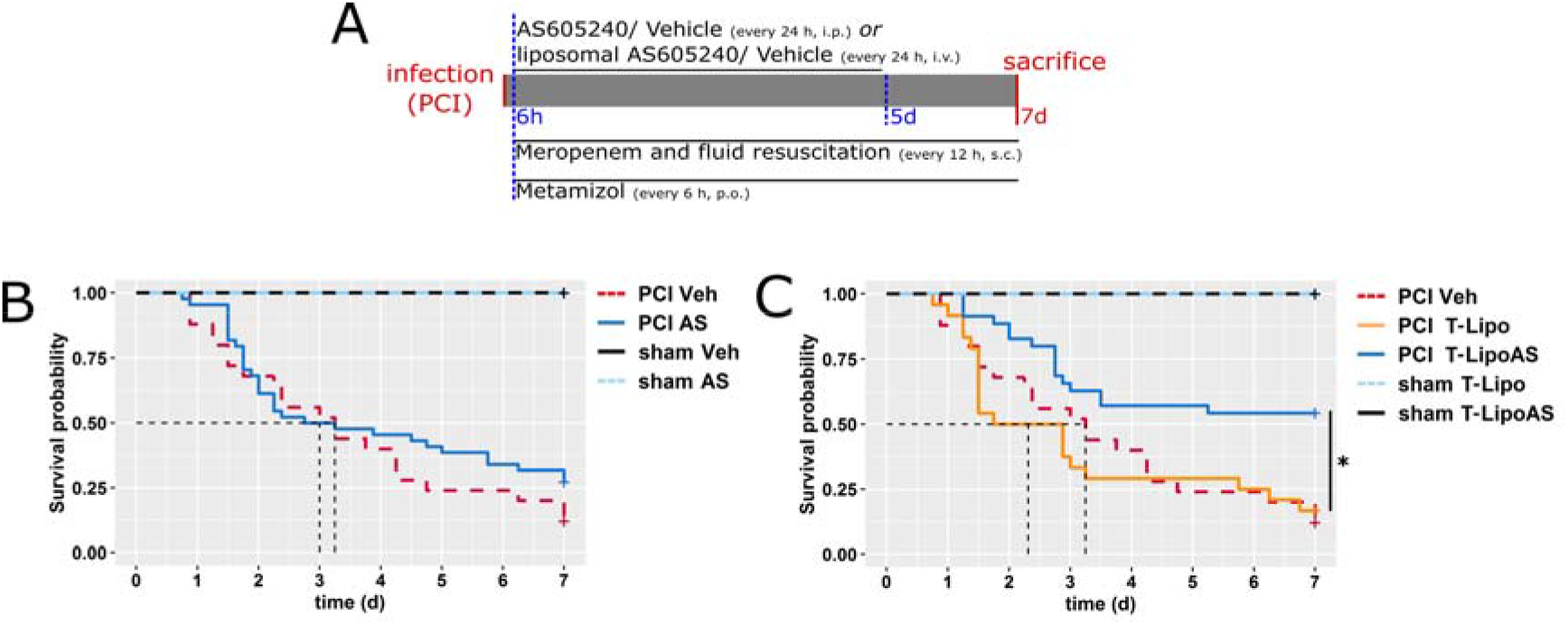
Restoration of excretory liver function in sepsis is associated with improved survival. (**A**) Treatment scheme and adjunctive therapy in the survival experiment. All animals received meropenem (25 mg kg^−1^ body weight, s.c.) twice daily and metamizole (2.5 mg per animal, p.o, every 6 h). (**B,C**) Survival analysis of PCI and sham animals. Subgroups were treated with (**B**) free AS605240 or vehicle (Veh; i.p.) or (**C**) with liposomal formulated AS 605240 (T-LipoAS) or T-Lipo (i.v.) as the specific vehicle control once per day. *p<0.05, Log-Rank Test

## Discussion

Sepsis afflicts approximately 1.7 million adults each year in the United States alone and contributes up to 270,000 in-hospital deaths.^27^ An improved understanding of clinical and biological interactions that can delineate disease phenotypes is key to identifying successful therapeutic approaches. Among the four clinical phenotypes described by Seymour and colleagues^28^ the phenotype characterized by impaired hepatic excretory function was associated with the worst outcomes. Experimental data identified down-regulation of canalicular transporters, especially Mrp2 as critical elements of impaired excretory function. Mechanistically, insertion of Mrp2 into ‘brush borders’ of the canalicular membrane has been found diminished and this phenotype depends critically upon PI3Kγ in sepsis models^4^. However, PI3Kγ is also recognized for its central role in leukocyte recruitment to sites of infection,^5,29^ a prerequisite for pathogen clearance, and thus a successful host response. Consistent with this notion, neither the ubiquitous knockdown of the PI3Kγ gene, nor its systemic pharmacological inhibition, improved survival despite the fact that the knockout prevents hyperbilirubinemia, the hallmark of cholestasis.^4^

To explore the potential separation of the cholestasis-preventing effect of PI3Kγ inhibition from untoward effects on immunity, we explored the cell-type-specific expression pattern of the gene in murine and human liver cells. We found that PI3Kγ is expressed across species in liver parenchymal cells, offering the possibility that selective inhibition of hepatocytes, if achieved, could improve liver function without impairing immune cell function.

Intriguingly, nanoformulation and targeting of the PI3Kγ inhibitor AS605240 allowed avoidance of untoward extrahepatic effects of PI3Kγ inhibition, for example on neutrophils, and improved outcomes in a murine model of sepsis. One can speculate why the effect of the kinase-inhibitor AS605240 on survival was more pronounced than the hepatocyte-specific PI3Kγ knockout; a maintained chaperoning function of the protein^30^ may be involved and would argue for a kinase-inhibitor rather than a gene silencing (siRNA) strategy.

Attempts to deliver small molecular drugs selectively to organs, e.g. hepatic parenchyma, thus holds considerable promise.^18^ This is in addition to well-developed and clinically available strategies for siRNA delivery^31^. Consistent with our previous results in PI3Kγ knockout mice^4^, systemic inhibition of PI3Kγ by intraperitoneally administered AS605240 led to an increased and prolonged cytokine storm compared to septic animals treated with the corresponding vehicle, indicative of impaired neutrophil migration and delayed resolution of infection. This cytokine storm was accompanied by a prolonged elevation of tissue damage markers, a significant fall in body weight and impaired clinical status reflecting significant morbidity. Thus, a protective effect of systemic PI3Kγ inhibition on hepatocellular excretory function appears to be antagonized by untoward side effects, including those acting on the immuno-inflammatory host response.

By contrast, septic animals treated with T-LipoAS showed no signs of an impaired immune response at the peritoneal site of infection. Thus, direct targeting of hepatocytes to prevent liver failure appears to preserve the host’s ability to fight infection. Neither PI3Kγ knockout nor systemic, non-targeted application of AS605240 could improve survival. Instead, non-targeted nanoformulations of AS605240 even led to an increase in mortality, possibly consistent with clearance of non-functionalized nanoparticles by innate immune cells thereby potentiating immunosuppressive off-target effects.

Local, inflammation-triggered alterations of the endothelial barrier, e.g. in a tumor micromilieu, likely contribute to the EPR effect that is used for targeted delivery of nanomedicines^10^. It is thus noteworthy that Dy635-targeting could achieve preferential accumulation of AS605240 within the liver while avoiding off-target effects on immunity, and this occurred even under conditions of systemic inflammation with presumably widespread altered barrier integrity.

As an obvious limitation, our study barely provides a proof of concept for a therapeutic intervention targeting parenchymal cells selectively. The feasible application of Dy635-specific targeting in a disease model beyond our previous report in healthy mice^12^ and the here documented expression of PI3Kγ in human hepatocytes should prompt, however, further translational studies.

## Conclusion

The present study demonstrates a druggable link between PI3Kγ activity and excretory liver failure in sepsis, a syndrome associated with poor outcomes. To overcome the off-target effects on immunity associated with systemic PI3Kγ inhibition, we successfully characterized a DY-635 conjugated liposomal carrier increasing the bioavailability of the PI3Kγ inhibitor AS605240 compartmentalized to hepatocytes, significantly improving liver function and survival. The simple and straightforward formulation as well as the described expression of PI3Kγ in human hepatocytes represents a significant opportunity for clinical translation of these carriers to target PI3Kγ-mediated liver failure with a simultaneous reduction of side effects on the immune system.

## Methods

### Liposomes

Liposomes were provided by BioLiTec Research GmbH, Jena, Germany. 1,2-Dipalmitoyl-sn-glycerol-3-phosphoethanolamine (DPPE) azide was conjugated to DY-635-alkin (Dyomics GmbH, Jena, Germany) and used to prepare liposomes using a 100 nm extruder. AS6052040-loaded liposomes were loaded with 0.2 mg AS605240 per 0.2 mg lipid mixture per mL in histidine buffer.

### AS605240 quantification

AS605240 concentration in liposomes was quantified by high-performance liquid chromatography coupled to tandem mass spectrometry (LC-MS/MS). The LC system (Shimadzu) was equipped with a CBM-20A Communication Bus Module, DGU-20A5R Degassing Unit, LC-20AD Prominence Liquid Chromatograph, SIL-20AC Prominence autosampler, and a CTO-20AC Prominence Column oven. A triple quadrupole mass spectrometer LCMS-8050 (Shimadzu) equipped with an APCI source operating in positive mode, was used for detection of AS605240 under the following parameters: nebulizing gas flow: 3 L min^−1^, heating gas flow: 10 L min^−1^, interface temperature: 350 °C, desolvation line temperature: 250 °C, heat block temperature: 200 °C, drying gas flow: 10 L min^−1^. Both quadrupoles, Q1 and Q3, were operated at a mass resolution medium with a dwell time of 9.0 ms for ergosterol and 6.0 ms for AS605240 (Abmole Bioscience). AS605240 was identified with an m/z of 258/187, and ergosterol with m/z 378/69 in multiple reaction monitoring mode. Gradient chromatographic separation was performed on a 2×;60 mm Multo-High100 RP-18 column (CS-Chromatographie Service, Langerwehe, Germany) with a particle size of 3 µm. The column was tempered at 50°C. Mobile phase A consisted of 0.1% (v/v) formic acid in deionized water and mobile phase B of 100% methanol. The column was equilibrated in 10% B with a flow rate of 0.4 mL min^−1^. The mobile phase was switched to 100% B after sample injection for elution. The flow rate was increased linearly from 0.4 mL min^−1^ at 3 min to 0.8 mL min^−1^ at 5 min and remained constant until 7 min. Subsequently, the mobile phase was changed to 10 % B in one step, and the flow rate was decreased linearly from 0.8 mL min^−1^ at 7.8 min to 0.3 mL min^−1^ at 8.5 min and remained constant until the end of the elution program at 9.5 min. Samples were diluted 1:10 in methanol, and 3 µM ergosterol (Sigma Aldrich, Germany) was added as an internal standard prior to detection. The injection volume of the sample was 3 µl. The analytical results were quantified with LabSolutions 5.91 (Shimadzu, Japan) based on an external AS605240 standard curve.

### DY-635 quantification

Liposomes were prepared as described for AS605240 quantification. DY-635 was detected using the Merck-Hitachi Elite LaChrom liquid chromatography (LC) system (VWR, Darmstadt, Germany). The LC system was equipped with the L-7250 autosampler, Merck peltier sample cooler for L-7250, L-2130 liquid chromatograph, the L-2300 column oven and the L-2485 fluorescence detector. Chromatographical separation was performed on a 60×2 mm MultoHigh 100 RP 18 column with 3 μm particle size (CS-Chromatographie Service, Langerwehe, Germany) at a constant temperature of 50 °C using the following gradient: The column was equilibrated in 10% B (methanol) and 90% A (0.1% (v/v) formic acid in ddH2O) with a flow rate of 0.4 mL min^−1^. At 0.1 min the mobile phase switched to 100% B and 0.5 mL min^−1^. The flow rate increased linearly from 0.5 mL min^−1^ at 3.5 min to 0.7 mL min^−1^ at 6.0 min and from 0.7 mL min^−1^ at 6.0 min to 0.8 mL min^−1^ at 10 min and remained constant until 13 min. Subsequently, the mobile phase changed to 10% B at 13.1 min, and the flow rate decreased linearly from 0.8 mL min^−1^ at 13.5 min to 0.5 mL min^−1^ at 15.0 min. Finally, the flow rate decreased to 0.4 mL min^−1^ at 15.5 min and remained constant until the end of the program at 16.0 min. DY-635 eluted from the column at 3.7 min and was detected using 595 nm excitation wavelengths and 670 nm emission wavelength. Injection volume was 10 µl. DY-635 concentrations were quantified using the EZChrom Elite 3.1.3 software based on an external calibration curve.

### Dynamic light scattering

The hydrodynamic radius of the liposomes was determined by dynamic light scattering (Nanosizer Nano ZS, Malvern Instruments, United Kingdom) in deionized water. Samples were equilibrated for 180 s before 3 × 30 runs (10 s/run) were carried out at 25 °C (laser wavelength, λ = 633 nm). Scattering counts were detected at an angle of 173° and intensity-weighted sizes and apparent distributions obtained from the standard cumulant analysis.

### Analytical ultracentrifugation

Analytical ultracentrifugation was performed as previously described^17^, using an Optima AUC analytical ultracentrifuge (Beckman Coulter, USA) with an An-50 Ti eight-hole rotor, and using double-sector epon centerpieces with a 12 mm optical solution path length. The cells were filled with 420 μL of liposomes diluted in histidine buffer, and with 440 μL of buffer in the reference sector. For analysis of the sedimentation velocity data, the ls-g*(s) model in Sedfit (version 15.01b) was used with confidence levels (F-ratios) set to 0.68. This model represents a least-squares boundary analysis under the assumption of non-diffusing species, and resolves the apparent differential distribution of sedimentation coefficients allowing for further conclusions on the samples (**Figures 1**).

### Cryo-transmission electron microscopy

Cryo-transmission electron microscopy (Cryo-TEM) was performed on an FEI Tecnai G² 20 system (ThermoFisher Scientific, USA) at an acceleration voltage of 200 kV. Samples were vitrified utilizing a Vitrobot Mark IV system. Samples were blotted (8.5 µL) on Quantifoil grids (R2/2) and plunge-frozen in liquid ethane (blot force equivalent −3 mm and a blotting time of 1 s). The diameters were analyzed from a minimum of 500 liposomes per group using FIJI. Geometric size (diameter) distributions were plotted as violin/boxplot using R GNU.^32^

### Ethical statement

All experimental procedures on animals were approved by the local government authority of Thuringia, Thüringer Landesverwaltungsamt, after evaluation and recommendation by an independent ethical advisory board, and carried out in accordance with the approved guidelines. FVB/N mice were housed under specific pathogen-free conditions in the animal facility of the Jena University Hospital. All experiments were performed on mixed populations of male and female mice. For sepsis experiments, the researchers inducing the initial insult were blinded to the distribution of the mice into the different treatment groups.

### Polymorphnuclear leukocyte isolation

A crude polymorphnuclear leukocyte (LEU) preparation was isolated from 10 mL human blood of healthy volunteers (heparinized) as previously described by Kuhns et al.^33^ In brief, the cell fraction was first separated from platelet-rich plasma by centrifugation (400 x g, 10 min, ambient temperature). The platelet-rich plasma was centrifuged again at 2000 x g, 10 min, and at ambient temperature to obtain the autologous serum used in a later step. The cell layer was transferred into a 50 mL conical tube which was filled with HBSS (no calcium, no magnesium) to a volume of 25 mL. 25 mL 3% dextran in HBSS was added and the closed tube then inverted five times and cells allowed to sediment for approximately 35 min. The RBC-free supernatant containing the leukocytes was transferred into a fresh tube, filled to 50 mL with HBSS and the leukocytes spun down at 400 x g for 10 min at ambient temperature. The cell pellet was then resuspended in ice-cold 1x RBC Lysis Buffer (420301, Biolegend) and left on ice for 5 min before cells were spun down. The Leukocyte pellet was taken up in 2 mL HBSS and overlaid to a discontinuous Percoll gradient (Bottom layer: 2 mL of 51% (v/v) Percoll, Middle layer: 2 mL 42% (v/v) Percoll, both prepared from Percoll (P4937, Sigma Aldrich) solution and autologous plasma). The isolation of polymorphonuclear leukocytes (LEU) was realized by centrifugation at 400 x g for 10 min and by retrieving the LEU from the bottom in 2 mL HBSS, and followed by washing once and centrifugation. The pellet was used for further analysis.

### Hepatocyte isolation, cultivation and stimulation

Primary murine hepatocytes were isolated from male and female FVB/N mice. Animals had been sacrificed by overdose with ketamine and xylazine. All procedures were carried out after the death of the animal was confirmed. The abdomen was disinfected, opened and the liver perfused through the portal vein with various buffers prewarmed at 40°C (roughly 38°C when reaching the liver tissue) at 8 mL min^−1^. First Krebs-Henseleit Buffer (Biochrom, Germany) containing 2 g L^−1^ glucose and 4 U heparin per mL was perfused for 5 min, followed by perfusion with Hepatocyte Digest Medium (HDM, ThermoFisher Scientific) for 10 to 15 min until the tissue was visually digested. Hepatocytes were strained through a mesh (70 µm, Corning) into 50 mL conical tubes and purified by centrifugation at 40 rcf, 4 min, 4°C 3 times. The hepatocytes in the pellet were kept and supernatant carefully replaced with Hepatocyte Wash Medium (ThermoFisher Scientific) in-between centrifugation steps. Primary human hepatocytes (huHEP) (Lonza, Swizerland) and isolated murine hepatocytes (muHEP) were seeded on rat-tail collagen (10 µg cm^−1^, Merck Millipore) coated well plates at a density of 0.1×10^6^ cells per cm². HuHEP were seeded in supplemented hepatocyte plating medium (HPM) (#MP100, Lonza, Swizerland) at 37°C. After 1 h the medium was carefully replaces with supplemented hepatocyte maintenance medium (HMM) (#MP250, Lonza, Swizerland) until stimulations. MuHEP were seeded and cultivated in Williams E Medium (PanBiotech, Germany) supplemented with stable glutamine (GlutaMaxx, ThermoFisher Scientific), insulin (Sigma Aldrich, Germany) and hydrocortisone (Sigma Aldrich, Germany), 10% fetal calf serum (ThermoFischer Scientific), and Penicillin/Streptomycin (ThermoFisher Scientific) (supplemented WEM).

After 16 h primary hepatocytes were stimulated with a mix of 100 ng mL^−1^ LPS (from *Escherichia Coli* O111:B4, #L2630, Sigma Aldrich), and species autologous cytokines: 10 ng mL^−1^ IFN-γ, 10 ng mL^−1^ IL-1β, and 50 ng mL^−1^ TNF-α (Prospec Biotech, Saudi Arabia) for 24 h in supplemented WEM (muHEP) or HMM (huHEP). Afterwards, cells were briefly washed in ice-cold PBS and lysed in RIPA buffer containing phosphatase inhibitor cocktail (PhosStop, Roche, USA) and protease inhibitor cocktail (HaltProtease Inhibitor Cocktail, ThermoFisher Scientific, USA).

### Uptake of liposomes in primary human hepatocytes

Transporter-qualified human hepatocytes from different donors were purchased from Lonza. Hepatocytes were seeded collagen-coated in 96 well plates using HPM and HMM as described above. The next day, after an equilibration period of approximately 8 h with fresh HMM, half of the cells were placed on ice for another 60 min. A two-fold concentrated ciclosporin A and liposome mixture (pre-tempered at 37°C or 4°C) was added to the same amount of culture medium to start the experiment. After 30 min another volume of ice-cold stop solution (10 mmol L-^1^ ciclosporin A) was added and cells spun down at 1000 rcf for 3 min. Cells were quickly washed once with HBSS by centrifugation and lysed in 10% ethanol in PBS by several freeze-thaw cycles. Solutions were then cleared by centrifugation at 4000 g for 10 min and the supernatant analyzed in a black 96 well microtiter plate using an EnSpire (PerkinElmer, USA) microplate reader with an Excitation/Emission wavelength of 630 nm/ 660 nm (with 9 nm band-passes around the maximum); against a standard in 5% ethanol, PBS. 10 µL of solution was then transferred to a clear 96 well plate and 0.2 mL BCA reagent (BCA Macro Kit, Serva Gel Electrophoresis GmbH) added. The protein concentration was determined using a bovine serum albumin standard curve in 10% ethanol/PBS. Absorbance was measured after 30 min incubation at 37°C at 562 nm. Background (cells not incubated with liposomes) was subtracted for all samples and dye concentrations normalized to the protein amount in each individual well. Experiments with hepatocytes of different donors were performed. Four individual wells for each condition were analyzed per run. Data were analyzed and plotted using v3.2.0 x64 together with the GRmetrics and ggplot2 plugin.^32,34^

### Assay of AS605240 activity *in vitro*

The activity of the PI3K inhibitor AS605240 (AbMole BioScience, USA) was assessed in primary human hepatocytes (Lonza, Swizerland) cultivated in collagen type I (10 µg cm^−1^) well plates for 16 h prior stimulation. The next day, cells were pretreated with 200 nmol L^−1^ AS605240 in 0.01% DMSO/PBS (w/o calcium and magnesium) or as a liposomal formulation for 1 h in HMM. Controls were pretreated either with either a 0.01% DMSO/PBS-solution. Cells were then stimulated for 20 min with 1 µmol L^−1^ fMLP (Sigma Aldrich) at 37 °C, washed with ice-cold PBS (w/o calcium and magnesium) and lysed immediately in RIPA-Buffer containing phosphatase inhibitor cocktail (PhosStop, Roche, USA) and protease inhibitor cocktail (HaltProtease Inhibitor Cocktail, ThermoFisher Scientific, USA). Protein content was quantified using the colorimetric bichinonic acid method (BCA Protein Assay Macro Kit, Serva Electrophoresis GmbH, Germany) and the same amount (25 µg) of protein loaded on an HSE Mini Gel (Serva Electrophoresis GmbH, Germany). Proteins were then transferred to 0.45 µm PVDF membranes (Carl Roth, Germany). Membranes are subsequently incubated in IgG-free 5% bovine serum albumin (BSA, CarlRoth, Germany) in tris-buffered saline containing 0.1% (v/v) Tween-20 (Carl Roth, Germany) (TTBS) for 1 h at ambient and stained using a Phospho-Akt (Ser473) (D9E) and Histone H3 (D1H2) (all Cell Signaling Technologies, UK) diluted 1:1000 in 5% BSA in TTBS. Membranes were developed after incubation with cross-adsorbed HRP-conjugated anti-rabbit antibody (#7047, Cell Signaling Technology) diluted 1:2000 in 5% BSA using ServaLight Eos CL or EosUltra CL HRP Substrate (Serva Electrophoresis GmbH, Germany) and a LAS ImageQuant 4000 (GE Healthcare, USA). Membranes were then stripped for 10 min in 200 mmol L^−1^ glycine, 1% (m/v) SDS and 1% Tween-20 at pH 2.2, washed in PBS and TTBS (2 times circa 10 min per buffer), blocked as described above and reprobed using a pan anti-Akt (C67E7) (#4691, Cell Signaling Technology) antibody (diluted 1:1000 in 5% BSA in TTBS) with the same detection system as described above.

The pAKT/AKT ratio was assessed by Western blot as an indicator for activity of the PI3Kγ and effects of the (liposomal) PI3Kγ inhibitor AS605240. Experiments were performed with hepatocytes from three different donors. The blots were analyzed densitometrically using ImageJ distribution Fiji v1.53g x64 Gel-analyzer plugin.^35,36^

### Detection of PI3Kγ in cell lysates

Purified human hepatocytes from 20 male or 20 female donors (HEP), an isolated non-parenchymal cell fraction (NPC) pooled from 3 male donors containing cholangiocytes, Kupffer cells and liver sinusoidal endothelial cells, and the purified crude polymorphonuclear leukocyte (LEU) fraction were lysed in RIPA buffer containing protease and phosphatase inhibitor (3 x 106 cells mL-1). HEP and NPC were purchased from Lonza, Swizerland. 5 µL of each lysate (HEP and LEU) and 15 µl NPC were used for the detection of PI3Kγ by Western blot. Alternatively, approximately 20 µg of protein from cultivated primary human or murine hepatocytes lysates were used.

The Western blot was performed as described above, and developed using ServaLight EoS CL Luminol and the LAS ImageQuant 4000 detection system (GE Healthcare, USA). The human PIK3CG (PI3Kγ) (#NBO2-15071, Novus Biologicals) was diluted 1:2000 in 5% BSA in TTBS and incubated over night at 4°C, rocking with the membrane.

### Immunohistochemistry

To investigate PI3Kγ expression in human liver tissue, biopsies from patients with various mild to severe liver diseases (**Supplementary Table 2**) were investigated. Paraffin-embedded liver tissue sections (4 µm thickness) mounted on glass slides are used for the following staining procedure. Tissue sections were deparaffinized and permeabilized using fresh citric acid buffer (10 mmol L^−1^ sodium citrate, 0.05% Tween 20, adjusted to pH 6.0 with 1 mol L^−1^ HCl) for 25 min. In TTBS rinsed slides were blocked for 10 min at room temperature using DAKO peroxidase blocking solution (#S200230, Dako), washed again in TTBS, blocked in donkey serum (Equitech-Bio, #SD30-0100), for 1 h at room temperature and incubated in primary antibody, rabbit polyclonal anti-PIK3CG (PI3Kγ) (NBP2-15071, Novus Biologicals) at 4°C overnight diluted 1:200 in antibody diluent (#S3022, Dako). Negative controls were incubated with antibody diluent only. Next day the slide were rinsed in TTBS, incubated in HRP-conjugated goat anti-rabbit IgG (ab214880, abcam) diluted 1:200 in antibody diluent for 2 h at room temperature, washed in TTBS and incubated in DAB Substrate solution (SignalStain DAB Substrat Kit, #8059, Cell Signaling Technology). After 10 min slides were carefully rinsed with deionized water and counterstained with hematoxylin. Washed and ethanol-fixed tissue sections were then mounted using Eukit (#03989, Sigma Aldrich). Images were acquired on an Olympus AX70 inverted microscope equipped with a AxioCam MRc5 (operated using AxioVision SE64 Rel. 4.9), a Halogen Lamp (U-PS, Olympus), a 20x/0.5, and 40x/0.75 semi-apochromatic objectives (Olympus).

### Survival and tissue sampling

PI3Kγ null (knockout)^5^ and liver specific PI3Kγ knockout mice (PI3Kγ flox^tg/tg^ x AlbCre^(tg)/tg^)^37^, as well as the background strain C57BL/6J were bred in the animal facility of the Jena University Hospital. Pharmaceutical inhibition of the PI3Kγ by AS605240 or liposomal formulations was carried out in FVB/N mice (8-12 weeks). All animals were either subjected to a peritoneal contamination and sepsis by injecting a characterized human stool suspension (PCI group)^23,24^ or saline (sham group) intraperitoneally. Our studies were designed to fulfill the minimum quality threshold in pre-clinical sepsis studies including antibiotic and analgesic treatment as well as fluid resuscitation.^38^ The dose of human stool suspension injected intraperitoneally was optimized for the C57BL/6J and FVB/N background aiming at a 7d-survival of 50% mortality (C57BL/6J) or 75% (FVB/N) in wild-type or untreated mice, respectively. At 6 h after infection antibiotics and fluid resuscitation (meropenem 2.5 mg per kg body weight dissolved in Ringer’s solution to a volume of 2.5 mg mL^−1^) were administered every 12 h subcutaneously for 7 days. Metamizole (oral, all experiments on FVB/N mice) or buprenorphine (subcutaneous, experiments on C57BL/6J and genetically engineered mice) was given for pain relief. DMSO-dissolved AS605240 (AS; 4 mg kg^−1^ body weight) or DMSO (Vehicle, Veh; 4 mg kg^−1^ body weight) solution was administered intraperitoneally. T-LipoAS, T-Lipo, NT-LipoAS or NT-Lipo were injected intravenously (injected liposomes 0.1 mg kg^−1^ body weight). Treatment with the different inhibitors was applied for the first 5 days beginning at 6 h after infection and repeated every 24 h. Under isoflurane anesthesia organs were harvested for analysis after 24 h or 48 h. The death occurred following a terminal cardiac puncture to collect EDTA plasma. Post-mortem lavage of the peritoneum was performed with 2 mL PBS containing 5 mmol L^−1^ EDTA.

### Neutrophil infiltration

Neutrophil infiltration in the peritoneal cavity was assessed by flow cytometry. Cells from 0.5 mL lavage were concentrated by centrifugation (1000 g, 10 min) and stained against CD45, CD11b, CD115, and Ly6-C. Identified neutrophil populations (CD45^pos.^, CD11b^pos.^, CD115^neg.^, Ly6-C^low/neg.^) were counted using a BD Accuri (BD Bioscience, USA) and flow software (v2, Perttu Terho, University of Turku, Finland).

### Intravital Microscopy

FVB/N mice with (PCI) and without (sham) sepsis were treated with (nanoformulated) inhibitors or vehicle solutions plus antibiotics and metamizole analgesia, as described above. Intravital microscopy was performed at 24 h post-infection. 1 h before surgery carprofen (5 mg kg^−1^ body weight, s.c.) was administered. Surgery and subsequent intravital microscopy were carried out under deep isoflurane anesthesia. Animals were kept warm using a heating pad or on a warmed (34°C), humidified microscopic chamber. The surgery required to perform intravital microscopy has been previously described.^39^ Liposomes and dyes were administered through a tail-vein catheter. DY-635 conjugated liposomes were administered to some animals. To investigate biodistribution during sepsis, CDFDA was injected to probe hepatocyte function. At the end of the experiment, propidium iodide (ThermoFisher Scientific, USA) and Hoechst 33342 (Sigma Aldrich, Germany) were injected for live/dead cell discrimination. Animals were sacrificed at the end of the experiment while remaining under anesthesia.

### Multispectral Optoacoustic Tomography

Animals were anesthetized and placed in the Multispectral Optoacoustic Tomography inVision 256-TF (iTheraMedical, Germany). A liver section was then focused in the tomograph. An optoacoustic data-set was acquired every 15 seconds for duration of 20 minutes. At each time point ten images per excitation wavelength were averaged from two different excitation wavelengths i.e. 800 nm and 900 nm for ICG and mouse anatomy respectively. A baseline of 2 minutes was acquired before injecting approximately 20 µg indocyanine green, ICG (Verdye, Diagnostic Green, USA) intravenously through a tail-vein catheter. The acquired images were pre-processed using software ViewMSOT, iTheraMedical GmbH (Release 3.8.1.04). The images were reconstructed by back-projection mode with a filter range of 50kHz to 6.5MHz-IR. After reconstruction, image analysis was carried out on the processed multispectral MSOT hyperstacks by using ImageJ distribution Fiji v1.49b x64^35,36^ and R v3.5.0 x64.^32^ The time-resolved, multispectral MSOT hyperstacks were first subdivided into two image stacks: one representing the signal from anatomical structures obtained by illumination with 900 nm laser wavelength, the other corresponding to the ICG signal obtained by illumination with 800 nm laser wavelength. Regions of interest (ROIs) were drawn manually around animal contours in the first cross-sectional time frame guided by the anatomical image stack. In order to compensate for motion artifacts, rigid image registration was performed using the Fiji plugin MultiStackReg v1.45 (Busse, Bard Science Downloads) which is based on a pyramid approach to subpixel intensity-based registration.^40^ To this end, the anatomical image stack was used to obtain transformation matrices for each time frame, which were then applied to the corresponding time frames of the ICG image stack. To adjust for intrinsic intensity variations between scans and to ensure inter-scan comparability, intensity values of the ICG image stack were z-transformed according to

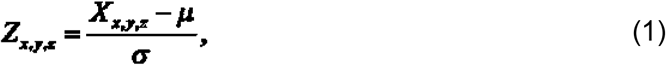

where ***Z***_***x,y,x***_ is the z-transformed intensity value at pixel position ***x,y*** in time frame ***z, X***_***x,y,z***_ is the original intensity value at pixel position ***x,y*** in time frame ***z***, and ***μ*** and ***σ*** are the mean and standard deviation of intensity values of the first-time frame. The time-resolved ICG image stack was then down sampled by 4 in the time domain to reduce the impact of animal breathing and was cropped to the bounding box of the ROI. All pixel values outside the ROI were set to zero. For each pixel in every ICG image stack, a vector of the signal intensity change rates between consecutive time frames was exported for further analysis. K-means clustering (k = 4) was applied to extract main characteristic time curves of signal intensity change rates for each treatment group (sham vehicle, sham T-LipoAS, PCI vehicle and PCI T-LipoAS). For each animal in a particular group, all pixel positions were compared to the characteristic curves that were obtained by k-means clustering for this treatment group. Each pixel position was assigned to belong to the characteristic curve that had the smallest Euclidean distance to the vector of signal intensity change rates at this pixel position. In that way, pixel abundance values were obtained for each animal with regard to the characteristic curves of the corresponding treatment group. To restrict the analysis to ICG uptake, only characteristic curves that represent a net increase of the ICG signal intensity over time were used for further quantification. The weighted average of those curves was calculated for each animal, using the pixel abundance values as weights and dividing by the sum of the respective pixel abundances. The tailing linear part of the thus resulting curves was then identified by visual inspection and found to start at time frame 8 (down-sampled time domain). Finally, the area under the curve was calculated for this linear part for each animal and used to compare treatment groups.

### Episcopic imaging of biodistribution

Mice injected with T-LipoAS were sacrificed and frozen post-mortem in TissueTek (OCT Compound, USA). An episcopic setup was applied on a cryomicrotome (CM3050 S Cryostat, Leica, Germany) for imaging of the whole mouse. DY-635 was excited with an LED at 625 nm wavelength (M625L4, Thorlabs, USA) and additionally filtered by a 1” bandpass filter with center wavelength of 630 nm and FWHM of 20 nm (ET630/20x, Chroma, USA). Fluorescence was observed through a 2” bandpass filter with a center wavelength of 670 nm and FWHM of 30 nm (ET670/30m, Chroma, USA). The sample was observed by a monochrome camera with 4500 x 3600 pixels and 6 µm pixel size (MicroLine ML16200, FLI Instruments, USA) equipped with a 1:1 lens with 89 mm focal length using f/4.8 (XENON-ZIRCONIA 2.8/89, Schneider Kreuznach, Germany). RGB images were generated by illuminating the sample sequentially with a ring of red, green and blue LEDs (24× SK6812RGBW-WS, Opsco, China).

### Plasma cytokine analysis

The Mouse Inflammation Panel 13-plex (LegendPlex, Biolegend, USA) was used to measure cytokine levels in EDTA plasma using filter plates. The samples were prepared, measured and analyzed using the BD Accuri according to the manufacturer’s instructions.

### Electron microscopy

For electron microscopy of the liver, animals were sacrificed at 24 h after the septic insult. The abdomen was opened and the liver perfused through the portal vein with buffer followed by a glutaraldehyde and formaldehyde based-fixative as described previously.^22^ Samples were washed with Dulbecco’s phosphate-buffered saline (DPBS, Sigma Aldrich, Germany) and subsequently processed by osmium tetroxide post-fixation (1% OsO_4_ in DPBS for 2 hours), and dehydration in a graded series of increasing ethanol:water mixtures (30%, 50%, 70%, 90%, 96% and 100%) for at least 10 minutes each. Embedding was performed by applying EPON812 embedding solutions (1:2 Epon: ethanol for 1 h, 1:1 for 2 h and 100% Epon overnight followed by a final exchange with 100% Epon and DMP). Samples were cured for 24 h at 60 °C. Samples were subsequently trimmed and sliced into 100-150 nm thick samples utilizing a PTPC Ultramicrotome (RMC Boeckeler, Germany) facilitated with a diamond knife (Diatome, Germany). Slices were placed on formvar/carbon-coated nickel finder grids (Plano, 200mesh). For improving the imaging contrast samples were stained with 2% uranyl acetate solutions for 30 minutes. Subsequently, all samples were analyzed by transmission electron microscopy (FEI Tecnai G² 20 at an acceleration voltage of 200 kV). Images were recorded utilizing an OSIS (Olympus Soft Imaging Solutions, USA) CCD camera with 1376 x 1024 pixels and analyzed by ImageJ software.

## Acknowledgement

We acknowledge U. Vetterling, S. Huschke, F. Haas, E. Jentho, F. Röstel, J. Guerra and J. Hoff (all Jena University Hospital, Germany) for their support in the animal experiments; Dr. F. Lehmann and Dr. M. Wenzel (both Dyomics GmbH, Germany) for spectral interpretation and chemistry; Dr. D. Steen, Dr. G. Wieland and Dr. D. Schegelmann on behalf of the Biolitec Research GmbH for producing and providing the various liposomal formulations and their generous technical support. Mr. A. Wiede (Leibniz-Institute for Photonic Technologies, Jena) and the mechanical workshop at the Jena University Hospital are acknowledged for their technical support and the construction of the episcopic imaging device.

## Contribution

**ATP, PB, JB** performed animal and cell culture experiments, imaging and tissue-analysis. **ATP, WF, IN, SH** and **TM** characterized nanomaterials. **MaBe** contributed to cell culture experiments. **BH** and **ZC** developed algorithms and analyzed MSOT data, **WH, WF** and **ATP** developed, performed and analyzed episcopic imaging. **SMC** and **MHG** supervised nanomaterial characterization. **NG** supervised experiments on human tissue sections. **RB** and **FG** performed the animal studies **RW** guided the study. **USS** supervised the material sciences concept. **MTF** supervised and guided MSOT image analysis. **ATP** and **MiBa** designed and guided the project. All authors contributed to writing the manuscript.

