## Supplementary Information for "Targeted delivery of a phosphoinositide 3-kinase γ inhibitor to restore organ function in sepsis through dye-functionalized lipid nanocarriers"

**Supplementary Table 1:** PI3Kγ expression in human tissue biopsies. LSECs: liver sinusoidal endothelial cells, KCs: Kupffer cells (local macrophages)

| **Sample information** | | **N** | **PI3Kγ expression** | | | |
|---|---|---|---|---|---|---|
| Diagnosis | Gender | | Hepatocytes | LSECs, KCs | Infiltrating immune cells | endothelial cells of larger vasculature |
| Non-alcoholic fatty liver disease (NASH) | female<br>male | 3<br>1 | yes | no | Yes | yes |
| Autoimmune hepatitis (AIH) | male<br>female | 1<br>1 | yes | no | Yes | yes |
| Liver cirrhosis | female<br>male | 2<br>1 | yes | no | Yes | yes |

**Supplementary Table 2:** Characterization of primary human hepatocyte donor pools (Lonza, Swizerland).

| Parameter | huHEP ♂$_{DP20}$ | huHEP ♀$_{DP20}$ |
|---|---|---|
| Number of Donors | 20 | 20 |
| Donor Age Average | 36.9 | 41.1 |
| Donor BMI Average[a] | 26.3 | 29.2 |
| Male Donors | 100.0% | 0.0% |
| Female Donors | 0.0% | 100.0% |
| Asian Donors | 5.0% | 0.0% |
| Afro American Donors | 5.0% | 10.0% |
| Caucasian Donors | 80% | 85.0% |
| Other Donors | 10.0% | 5.0% |
| No Drugs/ Alcohol/ Tobaco | 45.0% | 60.0% |
| Illicit Drug use | 35.0% | 20.0% |
| Tobacco Use | 35.0% | 20.0% |
| Heavy alcohol use[b] | 5.0% | 5.0% |
| Serologies CMV | Positive | Positive |
| Serologies EBV | Positive | Positive |
| Serologies HBV | Negative | Negative |
| Serologies HCV | Negative | Negative |
| Serologies HIV | Negative | Negative |
| CYP1A2 (pmol/ $10^6$ cells/min, 100 µmol $L^{-1}$ Phenacetin) | 68.7 | 81.7 |
| OATP1B1/3 Active Estrone-3-S Uptake (pmol/ $10^6$ cells/min, 10 µmol $L^{-1}$ Estrone-3-S)[c] | 1.6 | 1.9 |
| NTCP Active Taurocholate Uptake (pmol/ $10^6$ cells/min, 10 µmol $L^{-1}$ Taurocholate)[c] | 2.9 | 3.5 |

[a] The Body mass index (BMI) is derived body mass (weight) divided by the square of the body height of a person.

[b] Heavy Alcohol use is defined as >2 drinks per day or current history of alcoholism

[c] Transporter Activity is the amount of specific substrate retained on hepatocytes after incubation with 10 µmol $L^{-1}$ substrate at 4°C or at 37°C for 3 min. Active uptake is the fold change between uptake at 37°C versus 4°C. The uptake is expressed in pmol per $10^6$ cells per min.

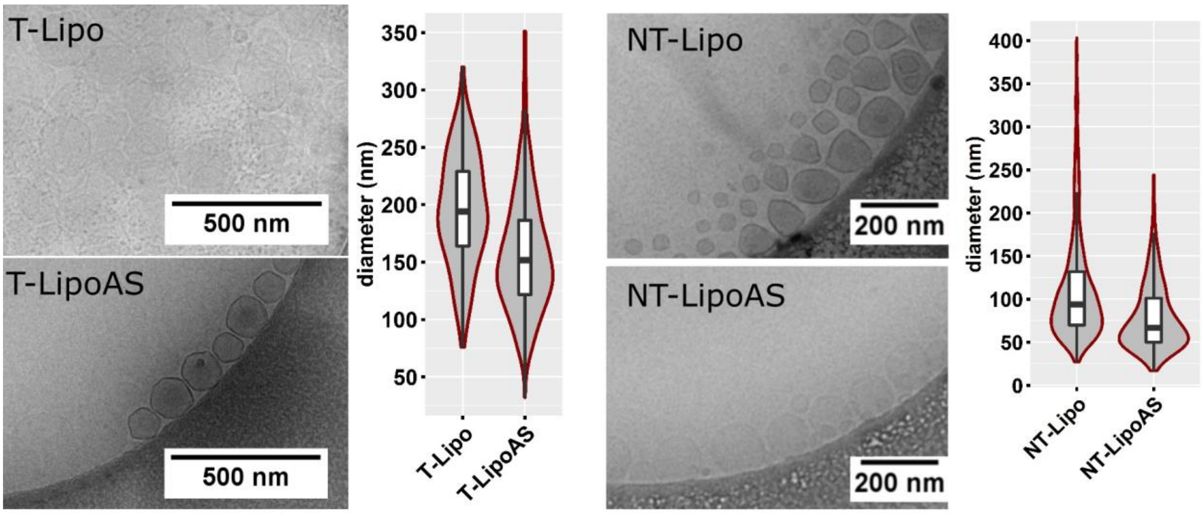


**Figure S1: Cryo-Transmission electron microscopy with size quantification of non-targeted Liposomal formulation.**

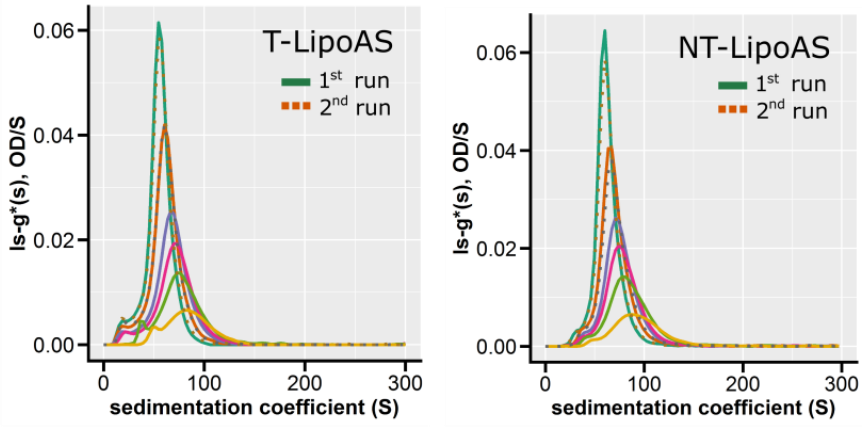


**Figure S2: Analytical ultracentrifugation of AS605240 loaded liposomes.**

Analytical ultracentrifugation was performed with different concentration of liposomes. The sedimentation was monitored by the optical density at 280 nm. Decrease of liposomal concentration shifted the sedimentation coefficient distributions toward larger values (solid lines). Sedimented liposomes were simply re-supsended by shaking of the ultracentrifuge cells, followed by another sedimentation velocity analysis (dotted lines). At all concentrations, very similar results were observed, indicating stability of drug-loaded liposomes during sedimentation and after resuspension leading to the observation of identical populations.

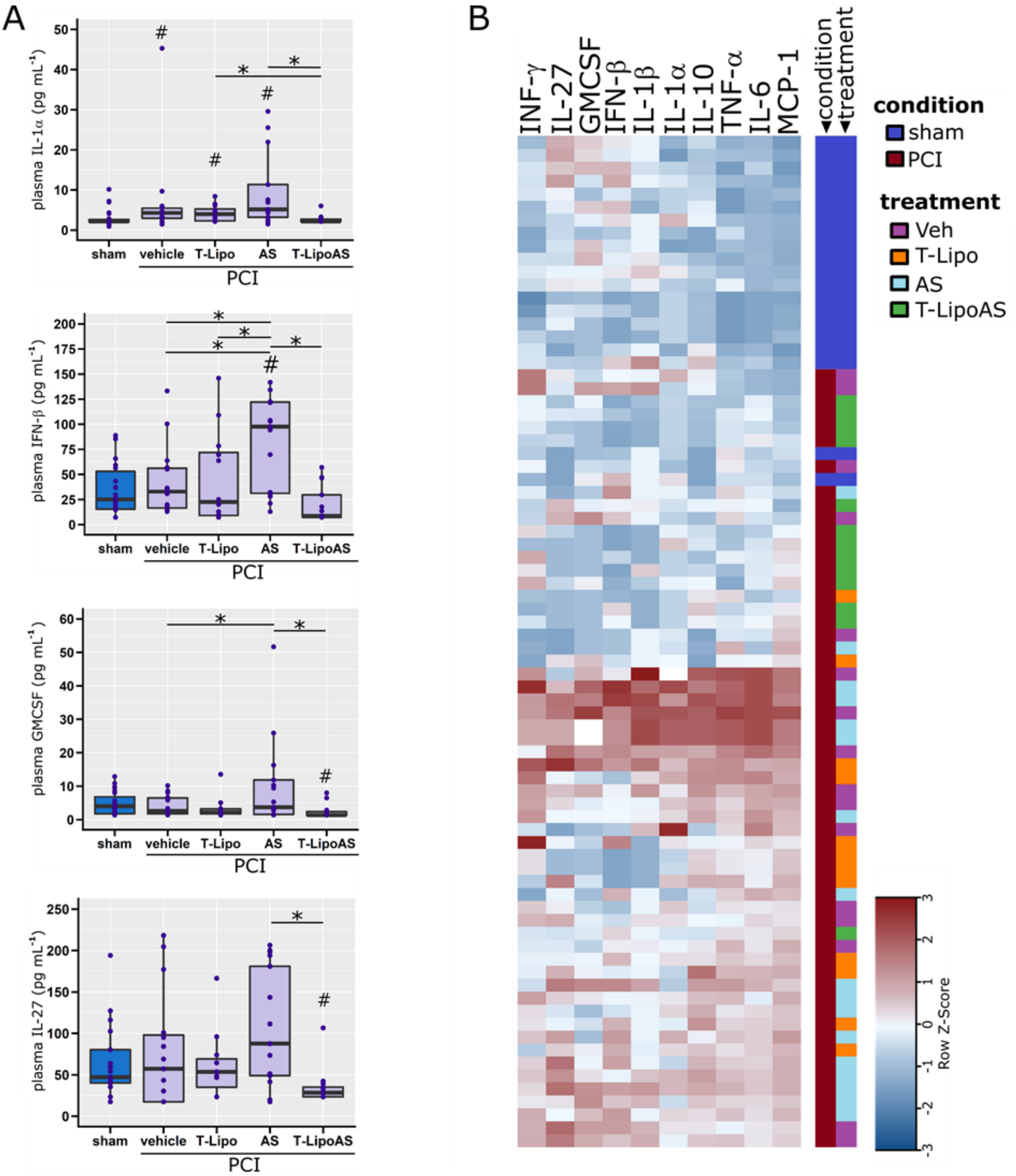


**Figure S3: Effects of targeted AS605240 delivery on the inflammatory response.**
**(A)** Pro-inflammatory-cytokines important for the host defense were analyzed in EDTA-plasma from PCI and sham animals. $^{\#}p<0.05$ against sham, $^{\#}*p<0.05$ against AS; Kruskal-Wallis ANOVA with controlled false-discovery rates (Benjamini-Hochberg procedure). **(B)** Cytokine profiles are depicted in a self-organizing heat-map according to their similarity of appearance in plasma.

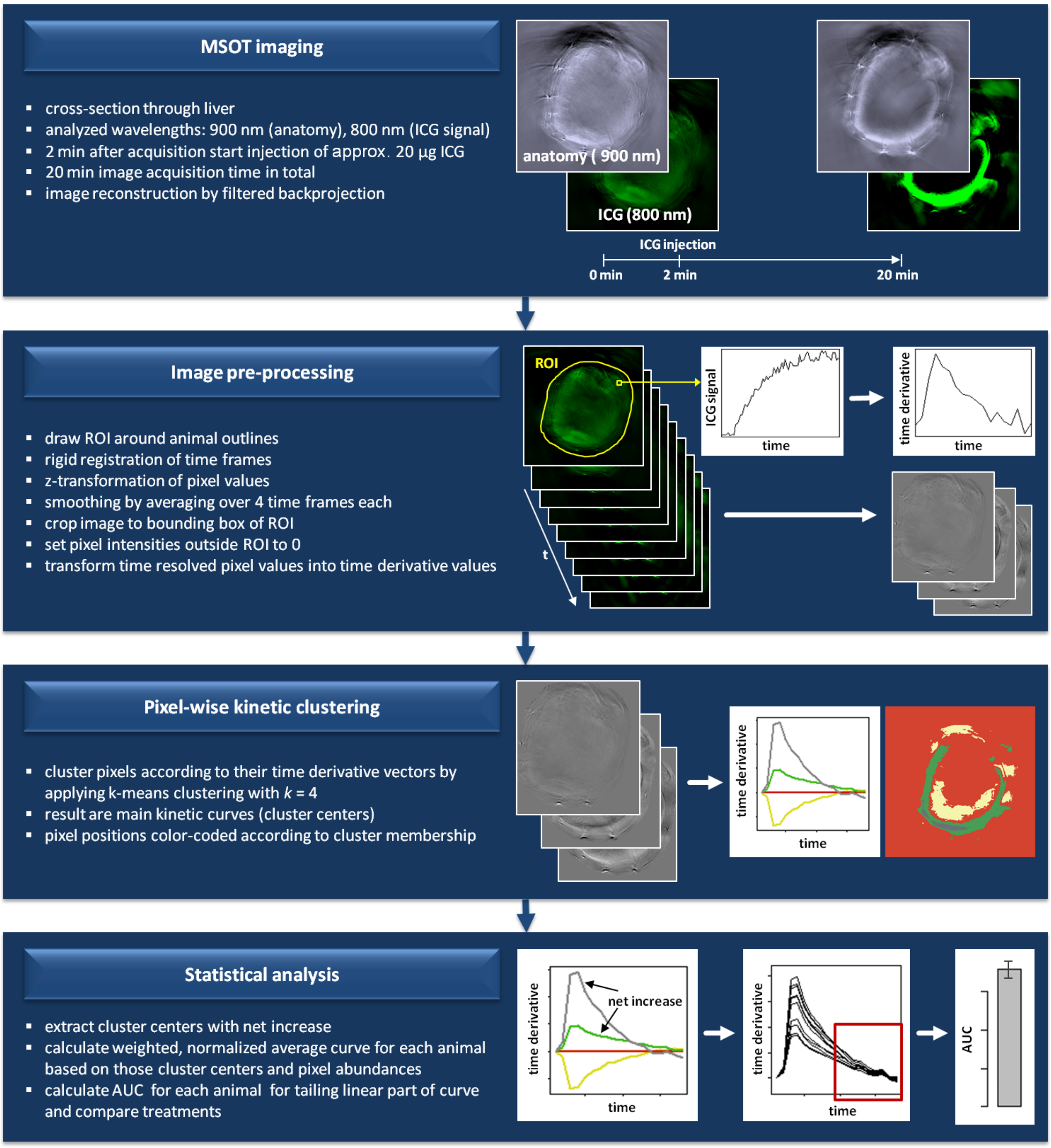


**Figure S4: Scheme on processing and analysis of multispectral optoacoustic tomography (MSOT) images for the quantification of liver function by indocyanine green (ICG) clearance.** MSOT imaging is performed to derive time-resolved cross-sectional images of liver tissue and ICG distribution. MSOT image stacks are first pre-processed to normalize ICG intensity values and smooth time course data. The time derivatives of pixel intensities within a region of interest (ROI) from one time frame to the next serve as input for pixel-wise kinetic clustering. By applying k-means clustering with k set to 4, 4 main kinetic curves (cluster centers) present in the ROI are extracted and color-coded images are generated to visualize the spatial distribution of those clusters for each animal. For quantification of ICG signal, only those cluster centers reflecting signal net increase are taken into account and combined into one curve, incorporating the respective abundances of pixels for each animal individually. All known three phases of ICG pharmacokinetics are clearly evident in those curves: first-pass (peak), re-distribution (exponential decay) and the final linear phase representing the dye-clearance through the liver. The area under the curve (AUC) was calculated from the linear decay phase to estimate the elimination capacity of the liver.

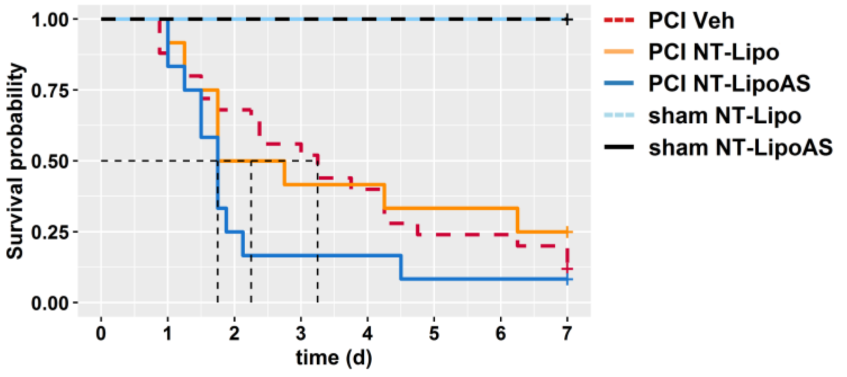


**Figure S5: Effects of the non-targeted liposomal AS605240 formulation on survival in the peritoneal contamination and infection (PCI) sepsis model.** Group sizes for sham groups NT-Lipo: 7, NT-LipoAS: 2 animals. For PCI (sepsis) groups: Veh: 25, NT-Lipo: 12, NT-LipoAS: 12 animals.The survival is monitored for seven days in this model. In the first two or three days the acute phase had passed and increasingly processes related to a cornification occur. Animals appear phenotypically healthy after five to seven days. However it is common in this model for abscesses to occur without visible symptoms, which in some cases may even be macroscopically detected as early as 24 h post infection, due to the lack of an surgical intervention cleaning and disinfecting the primary site of infection (here: abdominal cavity). These abscesses may grow and eventually rupture causing spontaneous death unrelated to the actual experiment. As a consequence the survival is limited here to seven days.
